# The oncogene FOXQ1 is a selective activator of Wnt/β-catenin signalling

**DOI:** 10.1101/2022.01.17.476620

**Authors:** Giulia Pizzolato, Lavanya Moparthi, Simon Söderholm, Claudio Cantù, Stefan Koch

## Abstract

The forkhead box transcription factor FOXQ1 is aberrantly induced in various cancers, and contributes to tumour growth and metastasis. It has been suggested that FOXQ1 exacerbates cancer by activating the oncogenic Wnt/β-catenin signalling pathway. However, the mode of action of FOXQ1 in the Wnt pathway remains to be resolved. Here, we report that FOXQ1 is a bimodal transcriptional activator of Wnt target gene expression in normal and cancer cells. Using co-immunoprecipitation, proximity proteomics, and reporter assays, we show that FOXQ1 engages the Wnt transcriptional complex to promote gene expression via TCF/LEF transcription factors. In parallel, FOXQ1 differentially regulates the expression of Wnt target genes independently of β-catenin and TCF/LEFs, which is facilitated by spatially separated activator and repressor domains. Our results suggest that FOXQ1 is a novel component of the Wnt transcriptional complex that reinforces and specifies Wnt signalling in a context-dependent manner.

## Introduction

The Wnt/β-catenin pathway is a major signalling cascade in development, tissue homeostasis, and stem cell maintenance (Clevers, 2006; MacDonald *et al*, 2009). Wnt pathway dysregulation frequently occurs in major diseases, notably cancer, in which activating pathway mutations aberrantly stabilise the transcription co-factor β-catenin (Nusse & Clevers, 2017). This allows β-catenin to enter the nucleus and activate T-cell factor/lymphoid enhancer-binding factor (TCF/LEF) family transcription factors, which drive a transcriptional program required for cell cycle progression and tissue self-renewal. Mounting evidence supports that the outcome of Wnt pathway activation is determined by numerous transcriptional co-regulators, which are recruited to discrete target genes in a tissue and context-specific manner (Söderholm & Cantù, 2020).

Forkhead box (FOX) transcription factors have emerged as one such family of Wnt pathway regulators (Koch, 2021). At least half of the 44 human FOX transcription factors are known to act as activators or inhibitors of Wnt signalling, but the function of most FOX proteins in the Wnt pathway is incompletely understood. Among these is FOXQ1, a known oncogene in several types of cancer (Bagati *et al*, 2017; Kaneda *et al*, 2010; Qiao *et al*, 2011). In colorectal cancers (CRC), in particular, FOXQ1 is one of the most highly upregulated genes, and has been linked to tumour growth and metastasis (Christensen *et al*, 2013; Kaneda *et al*., 2010). FOXQ1 has been identified as a candidate Wnt pathway activator in normal and CRC cells (Moparthi *et al*, 2019; Peng *et al*, 2015), and was shown to interact with β-catenin and TCF/LEF-associated Transducin-like enhancer (TLE) proteins (Bagati *et al*., 2017). It has been suggested that FOXQ1 activates Wnt signalling by promoting the nuclear translocation of β-catenin (Peng *et al*., 2015), or via the induction of canonical Wnt ligands (Xiang *et al*, 2020). However, the mode of action of FOXQ1 in Wnt signalling remains poorly defined.

Here, we identify FOXQ1 as a novel interactor of the TCF/LEF nuclear complex. We show that FOXQ1 reinforces TCF transcriptional activity in synergy with β-catenin. At the same time, FOXQ1 selectively controls the transcription of Wnt target genes in a β-catenin/TCF-independent manner. Consistently, we find that neither β-catenin binding nor Wnt ligand induction are required for the full activity of FOXQ1 in the Wnt pathway. Taken together, these observations suggest that FOXQ1 is a target-specific rheostat of Wnt transcriptional activity, which may have important implications for the pathobiology of colorectal cancer and the biology of FOX family transcription factor in general.

## Results

### FOXQ1 activates Wnt/β-catenin signalling in normal and colorectal cancer cells

Previous studies identified FOXQ1 as a candidate activator of Wnt/β-catenin signalling (Moparthi *et al*., 2019; Peng *et al*., 2015). We first confirmed these findings using a β-catenin/TCF luciferase reporter (TOPflash, (Veeman *et al*, 2003)) in normal and CRC cell lines (Fig 1A, B). In agreement with our earlier observations (Moparthi *et al*., 2019), we found that exogenous Flag-tagged FOXQ1 activated Wnt signalling in non-cancer 293T and HCT116 CRC cells, which harbour an activating β-catenin mutation and therefore have high basal Wnt activity (Fig 1B). Moreover, FOXQ1 strongly synergised with Wnt3a in TOPflash activation. We additionally observed that FOXQ1 differentially regulated several known Wnt targets, and significantly increased the expression of the prototypical target genes AXIN2 and SP5 (Fig 1C).

**Figure 1.**
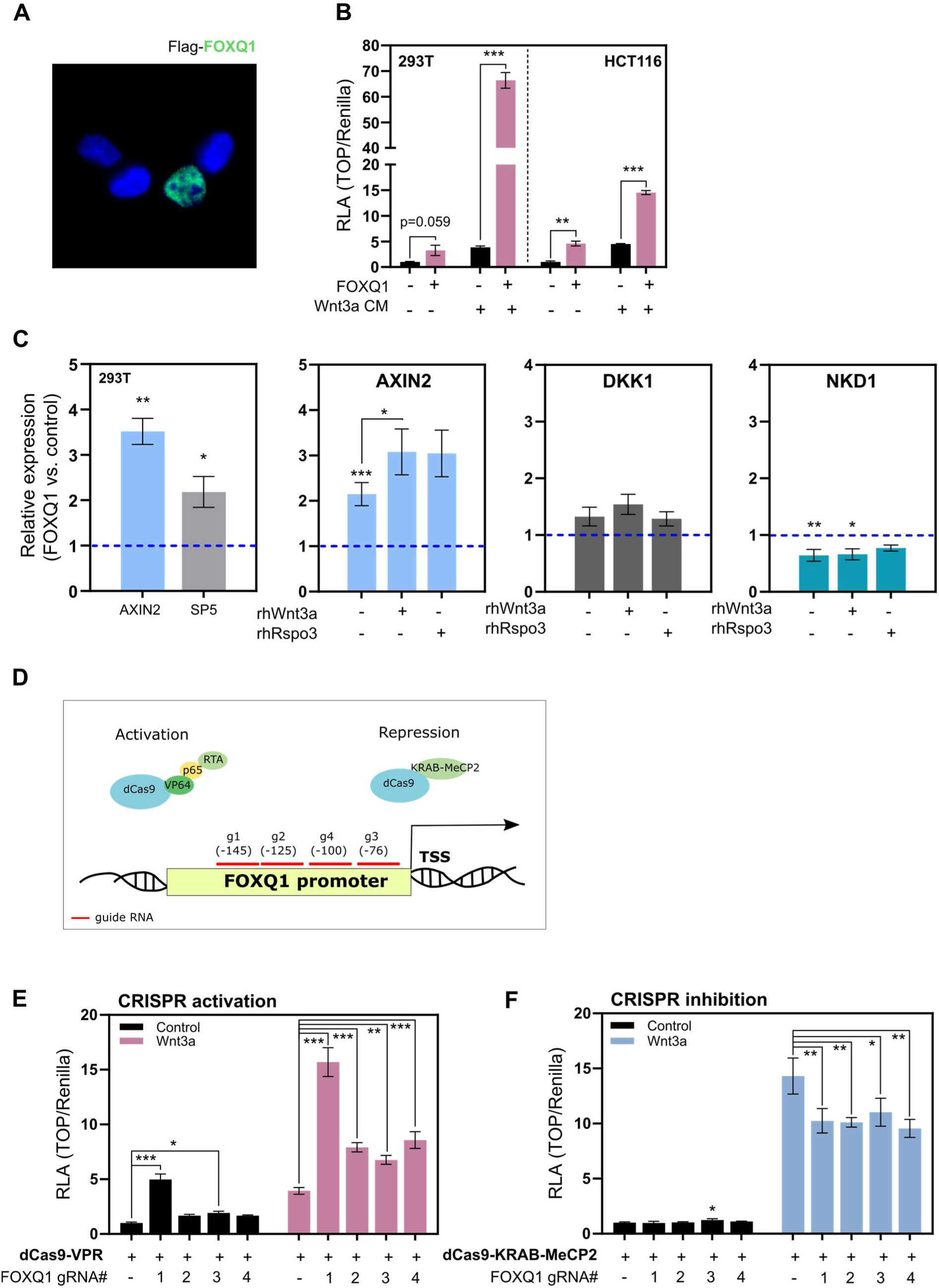
FOXQ1 activates Wnt/β-catenin signalling in normal and cancer cells. **A.** Representative immunofluorescence microscopy image showing nuclear localization of exogenous Flag-tagged FOXQ1 (green, with nuclei counterstained in blue) in colorectal cancer HCT116 cells. Original magnification: 40x **B.** FOXQ1 activates the β-catenin/TCF luciferase reporter TOPflash (TOP, normalised to Renilla control) in 293T and HCT116 cells, particularly in the presence of Wnt3a conditioned media (CM). Data were normalised to the untreated empty vector control for each cell line, and show one representative of n ≥ 3 independent experiments with biological triplicates. RLA, Relative Luciferase Activity. **C.** qPCR analysis of Wnt target gene expression in 293T cells. FOXQ1 induced *AXIN2* and *SP5* expression. Where indicated, cells were treated with recombinant human (rh) Wnt3a or R-spondin 3. *AXIN2* induction by FOXQ1 was significantly increased upon Wnt3a treatment. In contrast, FOXQ1 did not change *DKK1* expression and repressed *NKD1*. Samples were collected after 24 hours, and data from biological triplicates are displayed as fold change compared to empty vector control. **D.** Schematic representation of the CRISPR-mediated FOXQ1 transcriptional activation or inhibition (CRISPRa/i). dCas9-VPR or dCas9-KRAB-MeCP2 constructs were targeted to the FOXQ1 promoter using four non-overlapping guide RNAs (g1-4). Distance of gRNAs from the transcription start site (TSS) is indicated in parentheses. **E.** TOPflash reporter assay in 293T upon CRISPR activation of FOXQ1. Where indicated, cells were treated with Wnt3a conditioned media. *FOXQ1* induction by g1 and g3 significantly activated Wnt/β-catenin signalling in untreated cells. All gRNAs led to Wnt signalling activation in Wnt3a-treated cells. Data show one representative of n = 3 independent experiments with biological triplicates. **F.** TOPflash reporter assay in 293T upon CRISPR inhibition of FOXQ1. *FOXQ1* repression by g1-4 significantly reduced TOPflash activity after Wnt3a stimulation. Data show results from one experiment with biological triplicates. Data information: Data are displayed as mean ± SD. Statistical significance was determined by Welch’s t-test (B-C), or ANOVA with Dunnett’s post-hoc test (E-F), and defined as \**P* < 0.05, \*\**P* < 0.01, \*\*\**P* < 0.001.

To determine if physiological levels of FOXQ1 activate Wnt signalling, we next modulated FOXQ1 expression by CRISPR activation/inhibition (Chavez *et al*, 2015; Yeo *et al*, 2018). CRISPR activation using four non-overlapping guide RNAs targeting the FOXQ1 promoter increased *FOXQ1* levels in 293T cells up to 20-fold (Fig 1D, and Fig EV1A), which is a change in expression comparable to that observed in CRC (Christensen *et al*., 2013). Under these conditions, *FOXQ1* induction significantly increased TOPflash activity in 293T and HCT116 cells, especially in synergy with Wnt3a (Fig 1E, and Fig EV1B). Conversely, CRISPR inhibition of *FOXQ1* reduced Wnt reporter activity in Wnt3a-treated 293T cells (Fig 1F). We conclude that FOXQ1 is a physiologically relevant activator of Wnt/β-catenin signalling.

### FOXQ1 activates Wnt signalling downstream of β-catenin

To determine the mechanism by which FOXQ1 activates Wnt signalling, we performed epistasis assays in gene-edited 293T cells lacking essential pathway components. We observed a strong attenuation of Wnt activity in the absence of LRP6, and a complete loss of TOPflash activity in cells lacking LRP5/6 or β-catenin. (Fig 2A). This suggests that FOXQ1 cannot activate the Wnt reporter by itself, and that an intact Wnt receptor/β-catenin signalling axis is required for FOXQ1-dependent pathway activation. To support this conclusion, we first treated 293T and HCT116 cells with the porcupine inhibitor LGK974, which blocks Wnt ligand secretion (Fig 2B, C). TOPflash activation by FOXQ1 was inhibited by LGK974, and this reduction was particularly evident upon treatment with R-spondin 3, which increases Wnt receptor levels without affecting ligand abundance (Fig 2C, and Fig EV2A). In contrast, LGK974 had no effect in the presence of exogenous Wnt3a, and was less potent in HCT116 cells with constitutively stabilised β-catenin (Fig EV2A).

**Figure 2.**
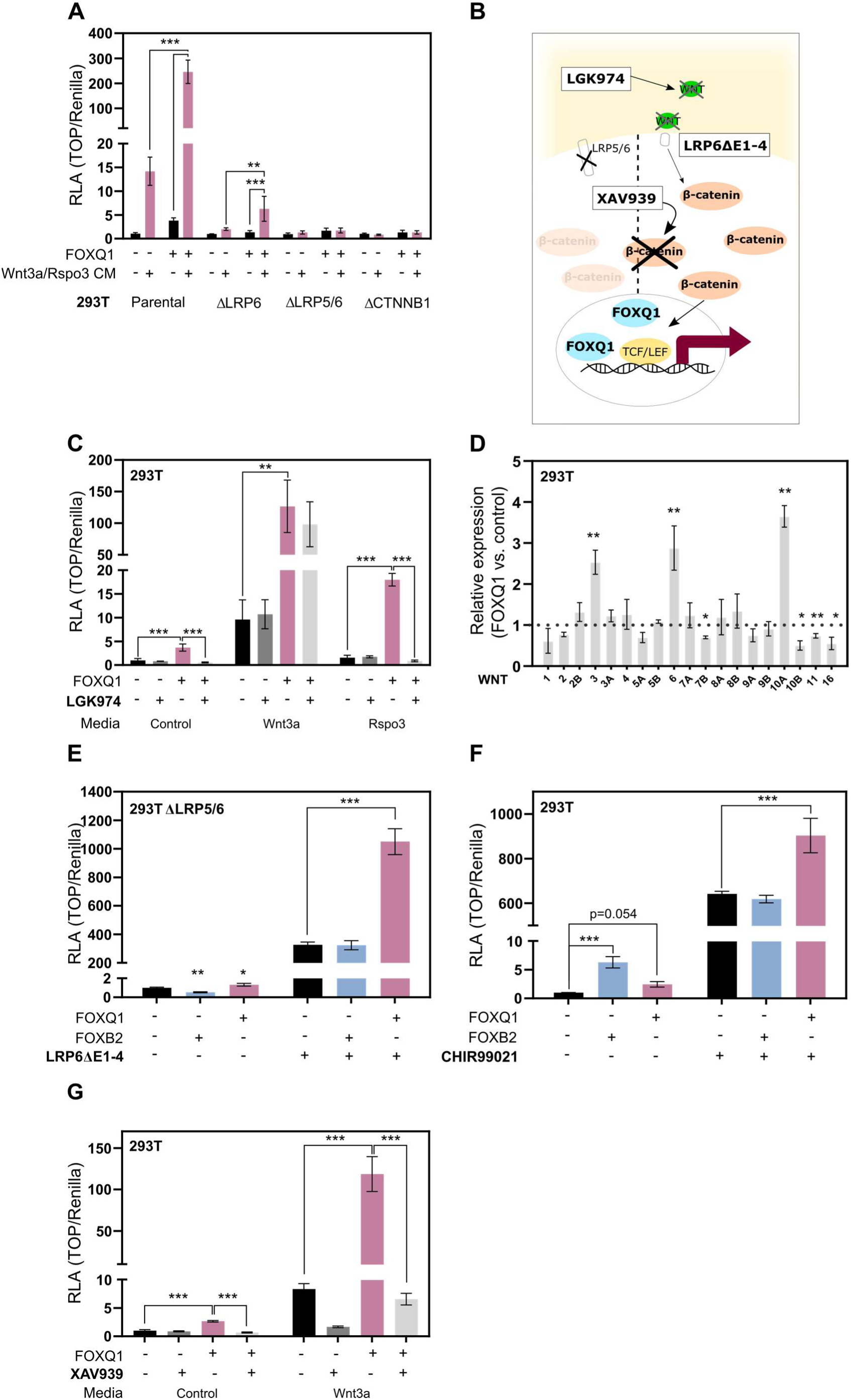
FOXQ1 activates Wnt/β-catenin signalling downstream of the receptor complex and β-catenin. **A.** Epistasis assay in normal and gene-edited 293T cells. Where indicated, cells were treated with Wnt3a and R-spondin 3 ΔC (Rspo3) conditioned media. Loss of the Wnt co-receptor LRP6 attenuated FOXQ1-dependent TOPflash reporter activation. Loss of LRP5/6 or β-catenin (CTNNB1) blocked reporter activation by FOXQ1. The graph shows one representative of n = 2 independent experiments with biological triplicates. **B.** Schematic showing the mode of action of the porcupine inhibitor LGK974, the constitutively active LRP6 ΔE1-4 construct, and the tankyrase inhibitor XAV939. **C.** TOPflash assay in 293T cells in the presence of LGK974. Treatment with LGK974 (10nM) attenuated FOXQ1-dependent Wnt activation, especially in the presence of exogenous Rspo3. Data show one representative of n = 3 independent experiments with biological triplicates. **D.** qPCR analysis of all 19 Wnt ligands in 293T cells. FOXQ1 significantly altered the expression of several Wnt genes, particularly *WNT3*, *WNT6*, and *WNT10A*. Samples were collected after 24 hours, and data are displayed as fold change compared to empty vector control from biological triplicates. **E.** TOPflash assay in 293T cells lacking Wnt co-receptors LRP5 and LRP6 (293T ΔLRP5/6). Where indicated, cells were transfected with a constitutively active LRP6 construct lacking the extracellular ligand binding domains (LRP6 ΔE1-4). FOXQ1, but not FOXB2, strongly activated the TOPflash reporter in the presence of LRP6 ΔE1-4. Data show one representative of n = 3 independent experiments with biological triplicates. **F.** TOPflash assay in 293T cells. Where indicated, cells were treated with GSK3 inhibitor CHIR99021 (5µM). FOXQ1, but not FOXB2, significantly activated the reporter construct in synergy with CHIR99021. Data show one representative of n = 3 independent experiments with biological triplicates. **G.** TOPflash assay in 293T cells. Where indicated, cells were treated with Wnt3a conditioned media and the tankyrase inhibitor XAV939 (5µM), which inhibits Wnt signalling by stabilising AXIN1. FOXQ1-dependent Wnt activity was significantly reduced upon β-catenin de-stabilization by XAV939. Data show one representative of n = 3 independent experiments with biological triplicates. Data information: Data are displayed as mean ± SD. Statistical significance was determined by ANOVA with Tukey’s (A, C, G) or Dunnett’s post-hoc test (E, F), or Welch’s t-test (D), and defined as * *P* < 0.05, \*\**P* < 0.01, \*\*\**P* < 0.001.

The aforementioned observations may be explained by induction of canonical Wnt ligands, as is the case for the similarly potent Wnt activator FOXB2 (Moparthi *et al*., 2019; Xiang *et al*., 2020). We thus examined the regulation of all 19 Wnt ligands by FOXQ1 in 293T cells. FOXQ1 significantly altered the expression of several Wnt ligands, particularly *WNT6* and *WNT10A* (Fig 2D, and Fig EV2B). In contrast to FOXB2 (Moparthi *et al*., 2019), however, expression changes were relatively modest. We therefore tested if FOXQ1-dependent Wnt ligand induction is required for its activity in the Wnt pathway. To this end, we uncoupled Wnt ligand binding from Wnt receptor activation by re-expressing constitutively active LRP6 (LRP6 ΔE1-4, (Davidson *et al*, 2005)) in 293T ΔLRP5/6 cells (Fig 2B, E). Neither FOXQ1 nor FOXB2 were able to activate TOPflash in LRP5/6-deficient cells, as expected. In contrast, FOXQ1 - but not FOXB2 - synergised with LRP6 ΔE1-4 in TOPflash activation (Fig 2E). Moreover, only FOXQ1 activated the Wnt reporter when β-catenin was artificially stabilised by the GSK3 inhibitor CHIR99021 (Fig 2F, and Fig EV2C). Conversely, we treated 293T and HCT116 cells with the tankyrase inhibitor XAV939, which decreases Wnt signalling by reducing β-catenin protein levels (Fig 2B, G). XAV939 significantly inhibited the activation of Wnt signalling by FOXQ1 (Fig 2G, and Fig EV2D). Finally, we determined TOPflash activity in 293T cells lacking β-catenin. FOXQ1 did not activate the reporter in the absence of β-catenin. However, FOXQ1 strongly synergized with exogenous wild-type or constitutively active (S33Y) β-catenin in these cells (Fig EV2E).

Taken together, these data suggest that FOXQ1 potentiates Wnt signalling downstream of β-catenin, irrespective of its ability to induce Wnt ligands.

### FOXQ1 interacts with components of the Wnt transcriptional complex

FOXQ1 interacts with β-catenin and TLE proteins (Bagati *et al*., 2017), suggesting that it may regulate TCF/LEF-dependent transcription. To test this hypothesis, we performed co-immunoprecipitation experiments in 293T cells. Consistent with previous observations (Bagati *et al*., 2017), overexpressed FOXQ1 precipitated both endogenous and exogenous β-catenin, as well as endogenous TLEs (Fig 3A, and Fig EV3A, B). Moreover, we observed that pull-down of TCF7L2 and LEF1 precipitated FOXQ1 in vitro, which was also the case in β-catenin-deficient cells (Fig 3B, and Fig EV3C). Some FOX proteins, such as FOXM1 and FOXG1, are thought to activate Wnt signalling by recruiting β-catenin to TCF/LEF (Zhang *et al*, 2011; Zheng *et al*, 2019). We therefore tested the association of TCF/LEF with endogenous β-catenin in nuclear extracts of Wnt3a-treated 293T cells. Compared to empty vector control, exogenous FOXQ1 increased β-catenin protein levels in the nucleus, and promoted a stronger association of β-catenin with TCF7L2 and LEF1 (Fig 3C, D).

**Figure 3.**
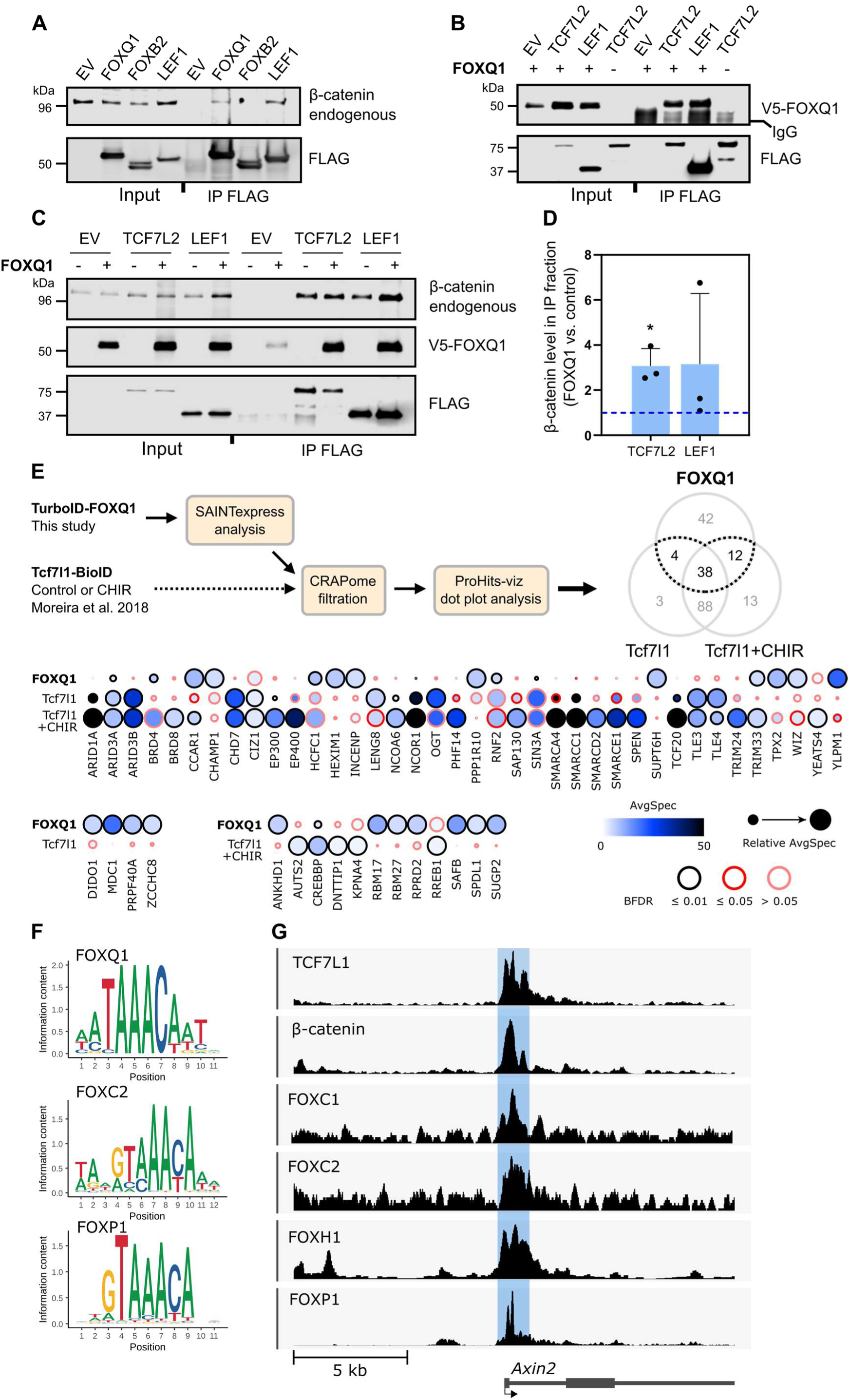
FOXQ1 interacts with the β-catenin/TCF/LEF transcriptional complex. **A.** Co-immunoprecipitation assay in nuclear lysates of 293T cells. Over-expressed Flag-tagged proteins were pulled down using a Flag antibody, and endogenous β-catenin was detected by immunoblotting. FOXB2 and LEF1 were used as negative and positive controls, respectively. **B.** Co-immunoprecipitation from nuclear lysates of 293T cells. Following pull-down of Flag-tagged TCF7L2 and LEF1, V5-FOXQ1 was detected by immunoblotting. **C.** Co-immunoprecipitation of Flag-tagged TCF7L2 and LEF1 from nuclear lysates of Wnt3a-treated 293T cells. Where indicated, cells were transfected with V5-FOXQ1. Following Flag pull-down, FOXQ1 and endogenous β-catenin were detected by immunoblotting. Data are representative of n = 3 independent experiments. **D.** Relative abundance of β-catenin associated with TCF7L2 and LEF1 from the previous immunoprecipitation experiments. FOXQ1 significantly increased β-catenin association with TCF7L2. **E.** Schematic representation of the workflow used for mass spectrometry data analysis. The Venn diagram highlights in black the number of shared protein between FOXQ1 and Tcf7l1 (untreated / CHIR99021-treated conditions). Dot plot analysis showing the 54 proteins that are common interactors of FOXQ1 and Tcf7l1. FOXQ1 experiments were performed with 4 biological replicates each for TurboID-FOXQ1 and control. AvgSpec, average spectral count; BFDR, Bayes false discovery rate. **F.** Sequence logo displaying the FOXQ1, FOXF2, FOXP1 consensus DNA-binding motif from the JASPAR database. **G.** Genomic tracks showing protein-DNA binding enrichment of TCF7L1, β-catenin, FOXC1, FOXC2, FOXH1 and FOXP1 at the *Axin2* locus, obtained by ChIP-seq. Data were retrieved from publicly available datasets (references and accession numbers in Figure EV4C). Data information: Data are displayed as mean ± SD. Statistical significance was determined by Welch’s t-test (D) and defined as * *P* < 0.05, \*\**P* < 0.01, \*\*\**P* < 0.001.

Our results so far suggested that FOXQ1 interacts with the β-catenin/TCF transcriptional complex. To expand on these observations and gain further insight into the function of FOXQ1 in the Wnt pathway, we generated a functionally active N-terminal TurboID-FOXQ1 fusion construct for proximity labelling proteomics (Branon *et al*, 2018; Cho *et al*, 2020), which we overexpressed in untreated 293T cells (Fig EV3D). Mass spectrometry analysis following streptavidin pull-down of biotinylated proteins identified nearly 400 candidate interactors that were significantly enriched compared to control. Gene ontology analysis of these hits revealed that FOXQ1 proximal proteins are primarily involved in mRNA processing, chromatin remodelling, and transcription regulation, but notably also β-catenin/TCF complex assembly (Fig EV3E, and Table 1). To identify common interactors of FOXQ1 and the Wnt transcriptional complex, we included a published BioID dataset of Tcf7l1 interactors in CHIR99021-treated or control mouse embryonic stem cells (Moreira *et al*, 2018) in our analyses. Following stringent, uniform data filtration and analysis, we observed that numerous candidate FOXQ1 interactors were shared with Tcf7l1 (Fig 3E, and Table 2). These included known regulators of Wnt/β-catenin signalling such as the histone acetyltransferases CREBBP/EP300 (Li *et al*, 2007), the chromatin remodelling factor SMARCA4 (Barker *et al*, 2001), and the transcription activator CCAR1 (Ou *et al*, 2009).

Consistent with a proximity of FOXQ1 to TCF/LEF, DNA sequence motif analysis identified several potential FOXQ1 binding sites within the chromatin regions where Wnt-responsive elements (WREs) are found (Fig EV4A). The DNA binding motif of FOX family transcription factors is highly conserved (Fig 3F, and Fig EV4B) (Dai *et al*, 2021). Thus, to determine whether these *in silico* predicted binding sites could be occupied by FOXQ1, we used an extensive series of available ChIP-seq datasets of other FOX proteins in varying cellular contexts as a proxy for potential FOXQ1 binding (Fig EV4C). Notably, many FOX transcription factors displayed physical occupancy of chromatin regions that are either overlapping (Fig 3G) or adjacent (Fig EV4D, E) to well described WREs at Wnt target gene loci.

Based on these observations, we conclude that FOXQ1 interacts with the TCF/LEF transcriptional complex, and that it promotes Wnt signalling by stabilising β-catenin/TCF interaction. Moreover, FOXQ1 may recruit and sequester additional transcription co-factors that increase the activity of β-catenin to WREs.

### Distinct protein domains shape transcription activation and repression by FOXQ1

To investigate the structural requirements for FOXQ1-dependent Wnt pathway regulation, we generated FOXQ1 constructs lacking either the N or C-terminus (Fig 4A, and Fig EV5A).

**Figure 4.**
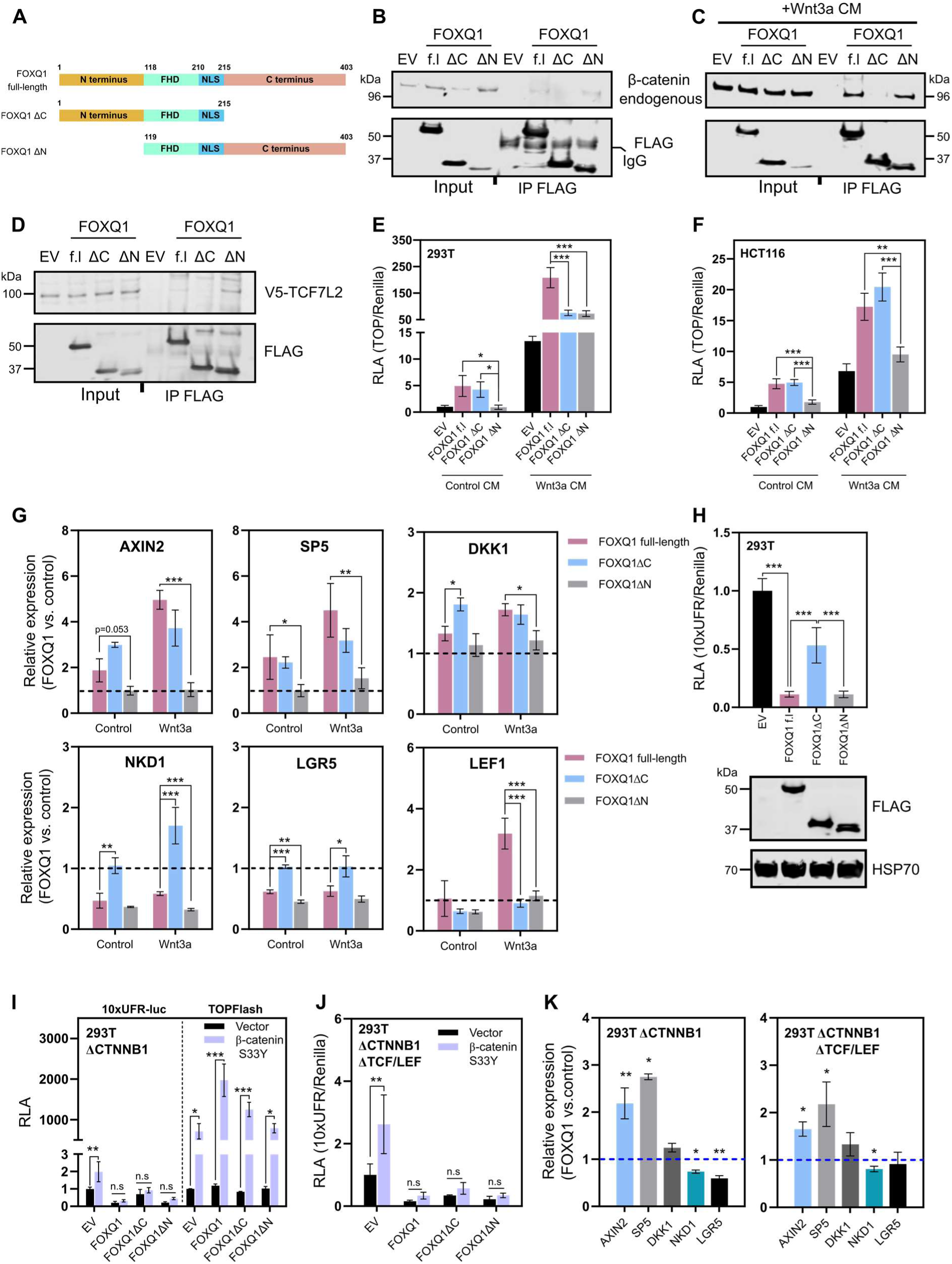
The FOXQ1 N and C-termini differentially regulate Wnt signalling. **A.** Schematic representation of Flag-FOXQ1 constructs used in subsequent assays. Numbers indicate amino acid positions. FHD: (DNA-binding) forkhead domain; NLS: nuclear localisation sequence. **B.** Co-immunoprecipitation assay in nuclear lysates of 293T cells. Following Flag pull-down of FOXQ1 constructs, endogenous β catenin was detected by immunoblotting. FOXQ1 ΔC was unable to bind β-catenin to any substantial degree. Representative blot from n = 2 independent experiments. **C.** Co-immunoprecipitation assay in nuclear lysates of 293T cells after treatment with Wnt3a conditioned media. As in the previous experiment, only full-length FOXQ1 and FOXQ1 ΔN interacted with β catenin. Representative blot from n = 2 independent experiments. **D.** Co-immunoprecipitation assay in nuclear lysates of 293T cells. Flag-tagged FOXQ1 constructs were pulled down in the presence of V5-TCF7L2. FOXQ1 ΔC was unable to bind TCF7L2 to any substantial degree. **E.** TOPflash assay in 293T cells. The FOXQ1 C-terminus was found to be dispensable for Wnt pathway activation in untreated (i.e., low Wnt) cells. In high Wnt conditions, neither construct activated TOPflash to the same extent as full-length FOXQ1. Data show one representative of n = 3 independent experiments with biological triplicates. **F.** TOPflash assay in HCT116 cells. The FOXQ1 ΔC construct activated Wnt signalling to the same extent as full-length FOXQ1. Data show one representative of n = 3 independent experiments with biological triplicates. **G.** qPCR analysis of selected Wnt target genes upon expression of FOXQ1 constructs in 293T cells. FOXQ1 ΔC induced *AXIN2*, *SP5*, and *DKK1* similarly to FOXQ1 full-length. In contrast, the FOXQ1 ΔN construct did not induce any Wnt target gene. *LEF1* was induced exclusively by full-length FOXQ1 upon Wnt3a treatment. **H.** FOXQ1 transcriptional activity upon expression of a forkhead box reporter plasmid (10xUFR-luc) in 293T cells. Loss of the FOXQ1 C-terminus resulted in significantly weaker transcriptional repression compared to the other constructs. Cell lysates from the assay were used for immunoblot to confirm equal expression. Data show one representative of n = 2 independent experiments with biological triplicates. **I.** Luciferase assay using the 10xUFR-luc and TOPflash reporters in 293T ΔCTNNB1 cells. FOXQ1 transcription repression at forkhead binding sites was not rescued by re-expression of β-catenin S33Y, despite their synergy in the TOPflash reporter assay. Data show one representative of n = 3 independent experiments with biological triplicates. **J.** Luciferase assay using the 10xUFR-luc reporter in 293T penta-knockout cells. β-catenin did not affect FOXQ1 transcriptional activity at forkhead binding sites in penta-knockout cells. Data show one representative of n = 3 independent experiments with biological triplicates. **K.** qPCR analysis of Wnt target genes in 293T ΔCTNNB1 and penta-knockout cells. FOXQ1 induced *AXIN2* and *SP5*, and repressed *NKD1* and *LGR5* in the absence of β-catenin and TCF/LEF proteins, albeit to a lesser extent as in parental cells. Data information: Data are displayed as mean ± SD. Where indicated, cells were treated with Wnt3a conditioned media. For qPCR experiments, samples were collected after 24 hours, and data from biological triplicates are displayed as fold change compared to empty vector control. Statistical significance was determined by Tukey’s post-hoc test following ANOVA (E-J) or Welch’s t-test (K), and defined as * *P* < 0.05, \*\**P* < 0.01, \*\*\**P* < 0.001 or n.s: not significant.

First, we performed co-immunoprecipitation experiments to identify the FOXQ1 domains that are required to engage the β-catenin/TCF transcriptional complex. In nuclear extracts of both untreated and Wnt3a-treated 293T cells, full-length FOXQ1 and FOXQ1 ΔN precipitated β-catenin (Fig 4B, C). Similarly, full-length FOXQ1 and FOXQ1 ΔN precipitated TCF7L2, whereas no interaction was detected with FOXQ1 ΔC (Fig 4D). In contrast, all constructs precipitated CREBBP, which was previously shown to mediate FOXP1-dependent Wnt pathway activation (Walker *et al*, 2015) (Fig EV5B).

Despite its inability to bind β-catenin and TCF, FOXQ1 ΔC promoted TOPflash activity in both control and Wnt3a-treated 293T cells, whereas FOXQ1 ΔN activated Wnt signalling only in the presence of Wnt3a (Fig 4E). Both deletion constructs exhibited reduced Wnt reporter activation in Wnt3a-treated 293T cells compared to the full-length protein, whereas the FOXQ1 C-terminus was completely dispensable for TOPflash activation in HCT116 cells (Fig 4F). We next examined the regulation of selected Wnt target genes by the FOXQ1 truncation constructs (Fig 4G). FOXQ1 ΔC induced *AXIN2*, *SP5*, and *DKK1* expression to a similar extent as full-length FOXQ1, and significantly derepressed *NKD1* and *LGR5* compared to full-length FOXQ1. In contrast, FOXQ1 ΔN was unable to activate Wnt target genes, but repressed *NKD1* and *LGR5*. Lastly, only full-length FOXQ1 induced *LEF1* expression, which required Wnt3a stimulation.

FOXQ1 acts as transcriptional repressor on its cognate forkhead box binding sites (Hoggatt *et al*, 2000; Moparthi & Koch, 2020). Thus, to generalise our findings, we additionally tested the FOXQ1 truncation constructs using the universal forkhead reporter 10x UFR-luc (Moparthi & Koch, 2020). Loss of the FOXQ1 C-terminus partially derepressed reporter activity, whereas loss of the N-terminus had no effect in this assay (Fig 4H). It has been suggested that β-catenin displaces TLE proteins from FOXQ1 to derepress *CDH2* (Bagati *et al*., 2017), which may explain the reduced Wnt responsiveness of the FOXQ1 ΔC construct (Fig 4G). However, re-expression of β-catenin in 293T ΔCTNNB1 cells or penta-knockout cells lacking β-catenin and all TCF/LEF proteins (Doumpas *et al*, 2019) had no effect on the regulation of 10x UFR-luc activity by any of the FOXQ1 constructs, despite their synergy in TOPflash assays (Fig 4I, J). Finally, we investigated the regulation of Wnt target genes in 293T ΔCTNNB1 and penta-knockout cells. FOXQ1 induced *AXIN2* and *SP5*, and repressed *NKD1* and *LGR5* expression in both cell lines, albeit to a lesser extent compared to wild-type cells (Fig 4K).

Collectively, these results suggest that FOXQ1 promotes β-catenin/TCF activity and represses specific target genes via N and C-terminal interactors, respectively. In addition, FOXQ1 acts as a selective transcriptional regulator of Wnt target genes in a β-catenin/TCF independent manner.

## Discussion

In this study, we identify the carcinoma oncogene FOXQ1 as a selective activator of Wnt/β-catenin signalling. Earlier studies suggested activation of Wnt/β-catenin signalling by FOXQ1 (Moparthi *et al*., 2019; Peng *et al*., 2015), but its mode of action in the Wnt pathway remained poorly defined. FOX transcription factors are known to control Wnt signalling through a variety of mechanisms, which include the induction of Wnt ligands, nuclear shuttling of β-catenin, and the stabilisation of Wnt transcriptional complexes (Koch, 2021). FOXQ1 appears to participate in many of these processes, consistent with earlier reports (Kaneda *et al*., 2010; Peng *et al*., 2015; Xiang *et al*., 2020), but our observations suggest that none of them can fully explain its activity in the Wnt pathway. Rather, we propose that FOXQ1 specifies the Wnt transcriptional output by acting both as an activator of β-catenin/TCF-mediated gene expression, and as an independent transcriptional regulator of Wnt target genes.

On the one hand, we find that FOXQ1 promotes the activity of TCF/LEF transcription factors, which strictly requires the presence of β-catenin. Although FOXQ1 physically interacts with β-catenin and TCF, this interaction is apparently largely dispensable for Wnt pathway activation. This observation, together with the fact that FOXQ1 increases Wnt reporter activity even at saturating β-catenin levels, suggests that it primarily affects the activity rather than the abundance or localisation of β-catenin. This resembles the proposed function of FOXP1, which was shown to activate Wnt signalling in B cell lymphoma (Walker *et al*., 2015). Walker and colleagues reported that FOXP1 recruits CREBBP to the Wnt transcriptional complex, which increases the acetylation and thereby the transcriptional activity of β-catenin (Li *et al*., 2007). We consider it likely that FOXQ1 recruits CREBBP and additional transcription co-factors such as EP300 via its N-terminus, which in combination activate β-catenin/TCF-dependent transcription. However, the validation and functional characterisation of FOXQ1 interactors will require further investigation.

On the other hand, we find that FOXQ1 differentially regulates the expression of specific Wnt target genes in a β-catenin/TCF-independent manner. The default mode of action of FOXQ1 is transcriptional repression (Hoggatt *et al*., 2000; Moparthi & Koch, 2020), which appears to be mediated by C-terminal interactors. Consistently, it has been shown that FOXQ1 negatively regulates the expression of *CDH2* by recruitment of TLE family repressors (Bagati *et al*., 2017), which may bind FOXQ1 via a C-terminal EH1 domain (Yaklichkin *et al*, 2007). Although we could not confirm general de-repression of FOXQ1-dependent transcription by β-catenin, it is possible that β-catenin cooperates with FOXQ1 in a gene-specific manner, as has been reported for class O FOX transcription factors (Doumpas *et al*., 2019; Essers *et al*, 2005; Tenbaum *et al*, 2012). It will therefore be of considerable interest to explore the genome-wide DNA binding pattern and transcriptome of FOXQ1.

FOXQ1 itself is a Wnt target gene (Christensen *et al*., 2013), and promotes CRC metastasis via induction of epithelial-to-mesenchymal transition (EMT) (Bagati *et al*., 2017; Kaneda *et al*., 2010; Qiao *et al*., 2011). Wnt pathway activation has been linked to EMT as well (Chen *et al*, 2012; Jiang *et al*, 2007; Nakayama *et al*, 2021), and FOXQ1 may thus act as a critical determinant of Wnt-induced EMT by balancing specific gene induction and repression in cancer cells. It is increasingly clear that a wide array of transcription factors stratifies Wnt/β-catenin signalling in cooperation with or in opposition to TCF/LEF (Bourgeois *et al*, 2021; Ramakrishnan *et al*, 2021; Söderholm & Cantù, 2020). Our results identify FOXQ1 as one such TCF-associated transcription factor, and highlight FOXQ1-dependent Wnt pathway regulation as a potential therapeutic vulnerability particularly in colorectal cancer.

## Materials and methods

### Plasmid / expression construct cloning

Molecular cloning of Flag/V5-tagged FOXQ1, FOXB2, and LEF1 has been described previously (Moparthi *et al*., 2019). Flag/V5-tagged TCF7L2 and Flag-tagged FOXQ1 truncation constructs were generated by restriction cloning using the high fidelity Q5 polymerase (New England Biolabs, Ipswich, US). For cloning of FOXQ1 truncation constructs, the following primers were used:

FOXQ1 ΔC- fw: CATGGAATTCAAGTTGGAGGTGTTCGTCCCTCG
FOXQ1 ΔC- rv: CATGCTCGAGTCAGCGCTTGCGGCGGCGGCG
FOXQ1 ΔN- fw: CATGGAATTCAAGCCCCCCTACTCGTACATC
FOXQ1 ΔN- rv: CATGCTCGAGGGCGCTACTCAGGCTAGGAGCGT

Flag-tagged TurboID was cloned to the N-terminus of FOXQ1. All plasmids were validated by partial sequencing (Eurofins Genomics, Ebersberg, Germany). Additional plasmids used in this study included Flag-LRP6ΔE1-4 (a gift from Christof Niehrs (Davidson *et al*., 2005)), mCherry-Beta-Catenin-20 (a gift from Michael Davidson; Addgene plasmid # 55001), and pcDNA3-S33Y Beta-catenin (a gift from Eric Fearon (Kolligs *et al*, 1999); Addgene plasmid # 19286).

### Cell culture and transfection

Authenticated 293T and HCT116 cells were obtained from the German Collection of Microorganisms and Cell Cultures (DSMZ, Braunschweig, Germany) and cultured in DMEM (Gibco) media supplemented with 10% fetal bovine serum (FBS), 2 mM glutamine and 1% (v/v) penicillin/streptomycin at 37 °C and 5% CO_2_. 293T ΔLRP6, 293T ΔCTNNB1, and 293T penta-knockout cells have been described previously (Doumpas *et al*., 2019; Moparthi *et al*., 2019).

293T ΔLRP5/6 cells were generated by transfecting 293T ΔLRP6 with an enhanced specificity Cas9 plasmid (eSpCas9(1.1), a gift from Feng Zhang (Slaymaker *et al*, 2016); Addgene plasmid # 71814) targeting LRP5 (gRNA: GGAAAACTGGAAGTCCACTG). Clonal cell lines were isolated by limiting dilution, and loss-of-function of LRP5/6 was validated by immunoblotting and functional assays, as before (Kirsch *et al*, 2017).

All cell lines were used at low passage and tested negative for Mycoplasma by analytical qPCR (Eurofins Genomics, Ebersberg, Germany). Wnt3a and control conditioned media were obtained from stably transfected L-cells, following the supplier’s guidelines (ATCC). R-spondin 3 conditioned media were generated by transient transfection of Rspo3 ΔC (Ohkawara *et al*, 2011) into 293T cells. Wnt3a and Rspo3 conditioned media were typically used at 1:4 and 1:1,000 dilution, respectively.

Cell transfection was performed using Lipofectamine 2000 (Thermo Fisher, Waltham, USA) or jetOPTIMUS transfection reagents (Polyplus Transfection, Illkirch, France), according to the supplier’s recommendations.

### Cas9-mediated transcription activation/inhibition

dCas9-VP64-p65-Rta and dCas9-KRAB-MeCP2 constructs for transcriptional programming (a gift from George Church (Chavez *et al*., 2015; Yeo *et al*., 2018); Addgene plasmid # 63798 / 110821) were used for CRISPRa/i. The constructs were guided to non-overlapping loci in the FOXQ1 promoter using four guide RNAs (gRNAs). The following oligo duplexes were cloned into the BPK1520 expression vector (a gift from Keith Joung (Kleinstiver *et al*, 2015); Addgene plasmid # 65777) to generate the gRNAs (g1-g4):

g1-fw: caccCCCAACGGGCGCGCACCAGG / g1-rv: aaacCCTGGTGCGCGCCCGTTGGG,
g2-fw: caccGCGCGCCCGTTGGGGAGCTG / g2-rv: aaacCAGCTCCCCAACGGGCGCGC,
g3-fw: caccGAGCGCGGACGGCAAGGGGT / g3-rv: aaacACCCCTTGCCGTCCGCGCTC,
g4-fw: caccCTGGGGAGCCGCCACCACCT / g4-rv: aaacAGGTGGTGGCGGCTCCCCAG

For validation of induction of FOXQ1 expression, cells were transfected in 24-well plates with 200 ng of dCas9-VPR and 10 ng gRNA in each well. RNA isolation was performed 48 hours after transfection. For reporter assays, cells were transfected in 96-well plates with 50 ng of dCas9-VPR or dCas9-KRAB-MeCP2, 5 ng gRNA, 50 ng of the TOPflash β-catenin/TCF reporter (M50 Super 8x TOPflash, a gift from Randall Moon (Veeman *et al*., 2003); Addgene plasmid # 12456) and 5 ng of Renilla luciferase control plasmid (a gift from David Bartel; Addgene plasmid # 12179) in each well. Cells were grown for 24 hours before luminescence measurement.

### Reporter assays

For the TOPflash assays, cells were seeded on a 96-well plate and transfected with 50 ng TOPflash reporter, 5 ng Renilla luciferase, and 10 ng of plasmid of interest in each well. Where indicated, 6 hours after transfection cells were treated with control, Wnt3a or Rspo3 conditioned media. For forkhead reporter assays, the TOPflash plasmid was replaced with 10x UFR-luc (Moparthi & Koch, 2020). The Dual luciferase assay was conducted as described previously (Hampf and Gossen, 2006) with few changes. Briefly, after overnight incubation, cells were lysed in passive lysis buffer (25 mM Tris, 2 mM DTT, 2 mM EDTA, 10% (v/v) glycerol, 1% (v/v) Triton X-100, (pH 7.8)) and agitated for 10 min. Lysates were transferred to flat bottomed 96-well luminescence assay plate. Firefly luciferase buffer (200 μM D-luciferin in 200 mM Tris-HCl, 15 mM MgSO4, 100 μM EDTA, 1 mM ATP, 25 mM DTT, pH 8.0) was added to each well and the plate was incubated for 2 min at room temperature. Luciferase activity was measured using Spark10 (Tecan) or a SpectraMax iD3 Multi-Mode Microplate Reader (Molecular Devices). Next, Renilla luciferase buffer (4 μM coelenterazine-h in 500 mM NaCl, 500 mM Na_2_SO_4_, 10 mM NaOAc, 15 mM EDTA, 25 mM sodium pyrophosphate, 50 μM phenyl-benzothiazole, pH 5.0) was added to the plate and luminescence was measured immediately. Data were normalized to the Renilla control values, performed in triplicate.

### Antibodies and reagents

The following antibodies were used: mouse anti-Flag M2 (F3165), rabbit anti-FLAG (F7425) from Sigma Aldrich (St. Louis, USA); rabbit anti-non phospho (Active) β-catenin (8814), rabbit anti-TLE1/2/3/4 (4681), rabbit anti-V5 (13202), rabbit anti-CBP (7389) from Cell Signaling Technology (Danvers, USA); mouse anti-FOXQ1 (C-9) (Santa Cruz sc-166265), mouse anti-V5 (ASJ-10004-100) from Nordic Biosite, rabbit anti-HSP70 from R&D Systems (AF1663).

Chemicals and inhibitors were from Sigma Aldrich and Cayman Chemicals (Ann Arbor, USA). CHIR99021, LGK974 and XAV939 have been characterized previously (Kulak *et al*, 2015; Liu *et al*, 2013; Naujok *et al*, 2014). Recombinant human WNT3A and R-spondin 3 were from R&D Systems (Minneapolis, USA).

### Immunocytochemistry

For FOXQ1 localization experiments, HCT116 cells were seeded on coverslips in 24-well plates. Cells were transfected with 200 ng of Flag-tagged FOXQ1 using Lipofectamine 2000 (Thermo Fisher, Waltham, USA). After 24 hours, cells were fixed with 4% paraformaldehyde, permeabilized with 0.1% Triton X-100 in PBS, and blocked with 2% bovine serum albumin (BSA) and 0.1% Tween-20 for 1 hour. Mouse anti-FOXQ1 was detected using fluorophore-labeled secondary antibodies (Thermo Fisher, Waltham, USA). Samples were mounted with Hoechst 33342 counterstain for nuclear visualization. Images were acquired on a Nikon E800 epifluorescence microscope (Amstelveen, Netherlands), and processed in ImageJ v1.52h (National Institutes of Health, Bethesda, USA).

### Quantitative real-time PCR

RNA extraction was performed using a Qiagen RNeasy mini kit (Hilden, Germany), and reverse transcribed (RT) with a Thermo Fisher cDNA synthesis kit. cDNA was amplified using validated custom primers, with SYBR green dye. Data were acquired on a Bio-Rad CFX96 Touch thermocycler (Hercules, USA), and normalized to *HPRT1* control. Data are displayed as fold change compared to empty vector control and show biological triplicates with technical duplicates.

### Immunoblotting and immunoprecipitation

Cells were harvested in PBS and lysed in lysis buffer (0.1-1% NP-40 in PBS with 1 x protease inhibitor cocktail). Lysates were boiled in Laemmli sample buffer with 50 mM DTT, separated on 10% polyacrylamide gels (Bio-Rad, Hercules, USA), transferred onto nitrocellulose membranes, and incubated in blocking buffer (LI-COR, Lincoln, USA). Primary antibodies were detected using near-infrared (NIR) fluorophore-labelled secondary antibodies (LI-COR, Lincoln, USA). Blots were scanned on a LI-COR CLx imager.

For co-immunoprecipitation experiments, cells seeded in 6-well plates and transfected with ≈1μg per well of the indicated constructs. Cells were harvested in PBS and nuclear extraction was performed using 0.1% NP-40 in PBS with 1x protease inhibitor cocktail. The proteins were pre-cleared using protein A/G agarose beads and immunoprecipitated by using anti-Flag M2 beads overnight at cold room. Samples were washed three times with 0.1% NP-40 in PBS, eluted in Laemmli buffer, and used for immunoblotting.

### TurboID and Mass spectrometry

The labelling and sample preparation of TurboID experiments was performed as described previously (Branon *et al*., 2018). Briefly, N-terminal TurboID-FOXQ1 and TurboID plasmids were transiently transfected into 293T cells using jetOPTIMUS (Polyplus Transfection, Illkirch, France). After 21 hours of transfection, cells were treated with 500 µM biotin, and incubated for 3 hours at 37 °C, 5% CO2. Cells were surface washed with ice cold PBS for three times to remove excess biotin and then harvested centrifuging at 1500 rpm for 15 min. Cells were washed thrice with ice-cold PBS buffer by centrifugation to remove any remaining biotin. Cells were lysed in RIPA buffer containing 1x protease inhibitor cocktail for 15 min on ice. Pre-washed streptavidin beads (GE Healthcare, USA) were added to the cell lysate and incubated overnight at 4 °C with end-over-end rotation. The beads were washed once with 1 ml of RIPA buffer, once with 1 mL of 1M KCl, once with 1 mL of 0.1 M Na2CO3, once with 1 mL of 2 M urea in 10 mM Tris-HCl (pH 8.0), and twice with 1 mL RIPA lysis buffer. The beads then transferred to new Eppendorf tube and washed twice with 50 mM Tris-HCl buffer (pH 7.5) and 2 M urea/50 mM Tris (pH 7.5) buffer. Beads were incubated with 0.4 μg trypsin (Thermo Fisher) in 2 M urea/50 mM Tris containing 1 mM DTT for 1 hour at 25 °C with end-over-end rotation. After incubation, the supernatant was collected and the beads were washed twice with 60 μL of 2 M urea/50 mM Tris buffer (pH 7.5) and the washes were combined with the collected supernatant. The supernatant was reduced with 4 mM DTT for 30 min at 25 °C with end-over-end rotation. The samples were alkylated with 10 mM iodoacetamide for 45 min in the dark at 25 °C with end-over-end rotation. For the complete digestion of the sample an additional 0.5 μg of trypsin was added and incubated at 25 °C overnight with end-over-end rotation. After overnight digestion the samples were desalted with C18 (thermos Scientific) Pipette tips and then dried with vacuum centrifuge.

TurboID samples were analysed by mass spectrometry, using an Easy nano LC 1200 system interfaced with a nanoEasy spray ion source (Thermo Scientific) connected Q Exactive HF Hybrid Quadrupole-Orbitrap Mass Spectrometer (Thermo Scientific). The peptides were loaded on a pre-column (Acclaim PepMap 100, 75um x2cm, Thermo Scientific) and the chromatographic separation was performed using an EASY-Spray C18 reversed-phase nano LC column (PepMap RSLC C18, 2um, 100A 75umx25cm, Thermo Scientific). The nanoLC was operating at 300 nL/min flow rate with a gradient (6-40 % in 95 min and 5 min hold at 100 %) solvent B (0.1% (v/v) formic acid in 100% acetonitrile) in solvent A (0.1% (v/v) formic acid in water) for 100 min.

Separated peptides were electrosprayed and analyzed using a Q-Exactive HF mass spectrometer (Thermo Scientific), operated in positive polarity in a data-dependent mode. Full scans were performed at 120,000 resolutions at a range of 380–1 400 m/z. The top 15 most intense multiple charged ions were isolated (1.2 m/z isolation window) and fragmented at a resolution of 30,000 with a dynamic exclusion of 30.0 s.

Raw data were processed by Proteome Discoverer1.4. (Thermo Fisher Scientific) searching the UniProt database with Sequest HT search engine. The search parameters were: Taxonomy: Homo sapiens; Enzymes; trypsin with two missed cleavages, no variable Modifications; fixed modification: Carbamidomethyl; Peptide Mass Tolerance, 10 ppm; MS/MS Fragment Tolerance, 0.02 Da. Quantification of the analysed data were performed with Scaffold 5.1.0, a Proteome Software using total spectral count. The mass spectrometry proteomics data have been deposited to the ProteomeXchange Consortium via the PRIDE partner repository with the dataset identifier PXD030464 and 10.6019/PXD030464.

### TurboID data analysis

Proteins indicated in the Proteome Discoverer output file as significantly increased in the TurboID-FOXQ1 samples (Fisher’s exact test < 0.05) were subjected to gene ontology analysis in DAVID v6.8 (Jiao *et al*, 2012), using the “BP Direct” function with default options. Processed mass spectrometry data were analysed further using SAINTexpress v3.6.3 (Teo *et al*, 2014). The resulting output file was merged with a dataset of Tcf7l1 interactors in mouse embryonic stem cells ((Moreira *et al*., 2018), their Supplemental Table S1), following mouse-to-human gene name conversion using the biomaRt R package (Durinck *et al*, 2009). Data were filtered against common mass spectrometry contaminants using the CRAPome repository (Mellacheruvu *et al*, 2013) with Frequency cut-off 0.2 or PSM ratio cut-off 3. Then, data were analysed and visualised in ProHits-viz (Knight *et al*, 2017), using the Dot plot analysis tool with default options.

### Forkhead box phylogenetic analysis

FOX transcription factor phylogenetic tree was constructed using the Molecular Evolutionary Genetics Analysis (MEGA) software (version 11) (Tamura *et al*, 2021). Forkhead box domain peptide sequences for each FOX transcription factor were downloaded from the UniProt database (UniProt Consortium, 2021) (accessed 2021-12-14). Multiple sequence alignment was performed using the ClustalW algorithm with default settings. Phylogenetic analysis and construction of Maximum Likelihood Phylogenetic Tree was done with default settings.

### External Chip-seq data

We performed a systematic review of publicly available FOX transcription factor ChIP-seq data from mouse. Datasets were downloaded from the Gene Expression Omnibus (GEO) database. Data based on older versions of the mouse reference genome were converted to version mm10 using the UCSC liftOver tool. Data files were further converted to BigWig file format before visualization in the Integrative Genomics Viewer (IGV) (Robinson *et al*, 2011).

### *In silico* FOXQ1 binding prediction

FOXQ1 transcription factor binding profile data was downloaded from the JASPAR database, 9^th^ release (2022) (Castro-Mondragon *et al*, 2021) as position frequency matrices (PFMs). Based on these binding profiles, FOXQ1 binding was predicted at the *Axin2* and *Lef1* loci (mouse reference genome mm10) using the R package TFBStools (Tan & Lenhard, 2016). Genomic regions for which to scan for FOXQ1 binding patterns were defined as to include sites previously identified as Wnt-responsive elements (WRE) (Jho *et al*, 2002; Li *et al*, 2006).

### Statistical analyses

Data are shown as mean with standard deviation (SD), as indicated in the figure legends. Each experiment included controls (e.g., empty backbone plasmid / substance carriers) at identical concentrations.

Where indicated, Welch’s *t*-tests (two groups) or one-way ANOVA analyses with Dunnett or Tukey post hoc tests (three or more groups) were calculated using R 4.1.1. Significance is indicated as: \**P* < 0.05, \*\**P* < 0.01, \*\*\**P* < 0.001.

## Data availability

The mass spectrometry proteomics data have been deposited to the ProteomeXchange Consortium via the PRIDE partner repository with the dataset identifier PXD030464 and 10.6019/PXD030464. All raw data are available upon request to the corresponding author.

## Acknowledgments

The authors thank all investigators who have made data and materials available to the scientific community. Technical support from the mass spectrometry core facility at Linköping University is gratefully acknowledged.

C.C. and S.K. are Wallenberg Molecular Medicine Fellows, and receive financial support from the Knut and Alice Wallenberg Foundation. Additional financial support came from project grants from the Swedish Research Council (VR), and the Swedish Cancer Society (Cancerfonden) to S.K.

## Author contributions

Conceptualization and Data curation - S.K. and L.M.; Formal Analysis - S.K., G.P., L.M. and S.S.; Funding acquisition, Resources and Supervision - S.K. and C.C.; Investigation and Methodology - G.P., L.M. and S.S.; Visualisation - G.P. and S.S.; Writing – original draft - G.P.; Writing – review & editing – S.K., L.M., S.S. and C.C.

## Supplementary Figures

**Figure EV1.**
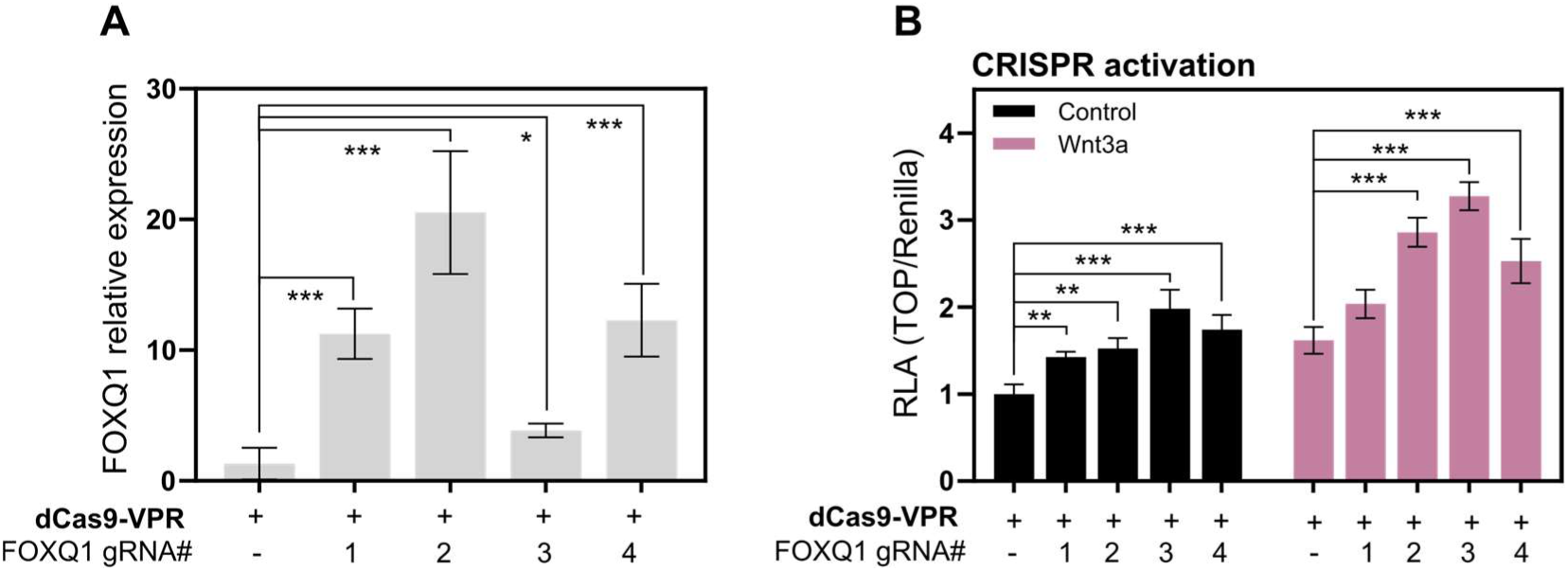
FOXQ1 is a physiological activator of Wnt/β-catenin signalling. **A.** *FOXQ1* qPCR in 293T cells. *FOXQ1* expression was significantly induced by a dCas9-VPR (CRISPR activation) construct guided to the *FOXQ1* promoter by 4 non-overlapping guide RNAs. Data are displayed as fold change compared to empty vector control and show biological triplicates. **B.** TOPflash reporter assay showing that CRISPR-mediated induction of FOXQ1 activated Wnt/β-catenin signalling in HCT116 cells. Data show one representative of n = 2 independent experiments with biological triplicates. Data information: Data are displayed as mean ± SD. Statistical significance was determined by ANOVA with Dunnett’s post-hoc test, and defined as * *P* < 0.05, \*\**P* < 0.01, \*\*\**P* < 0.001.

**Figure EV2.**
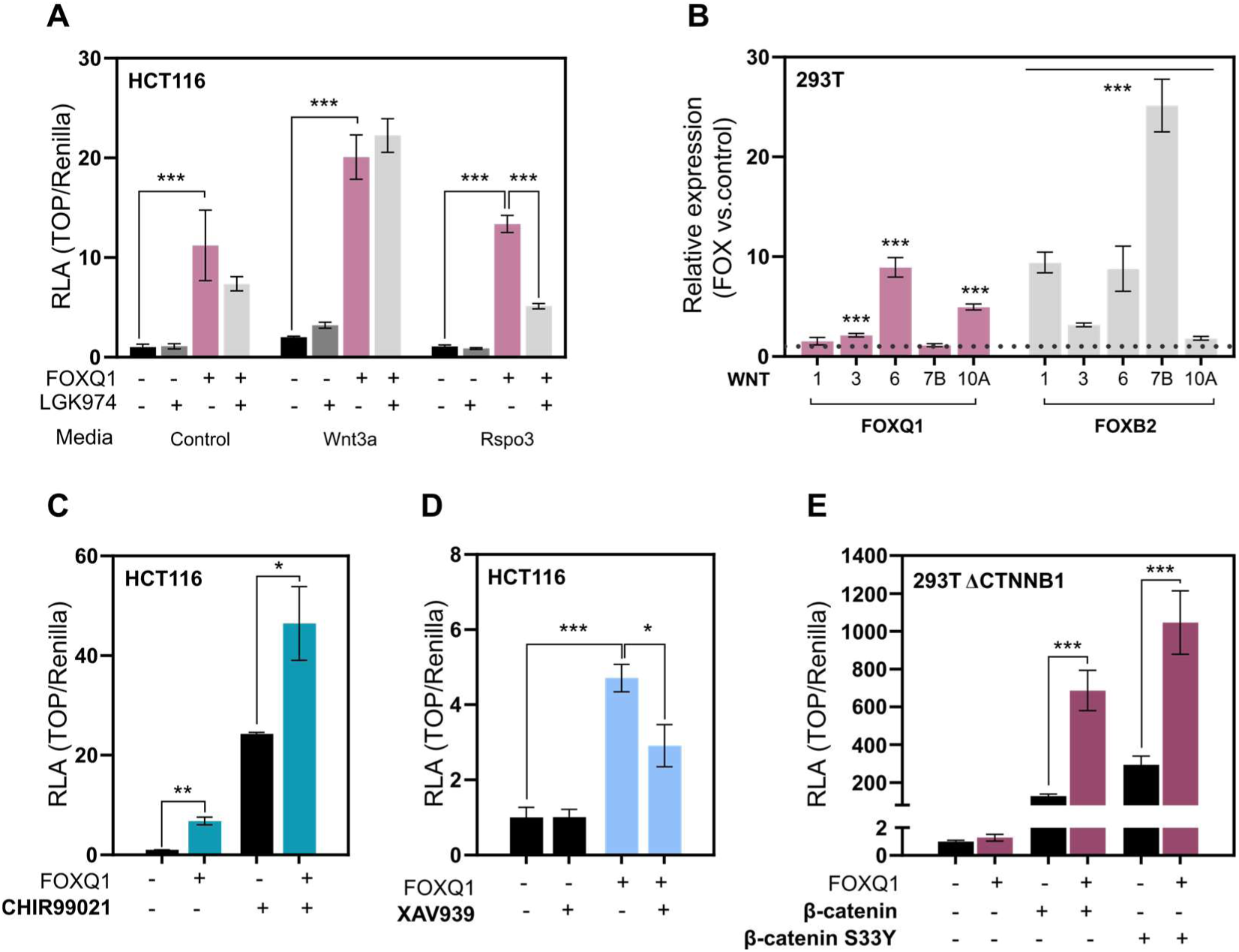
FOXQ1 requires upstream pathway activation and β-catenin stabilization to activate Wnt/β-catenin signalling. **A.** TOPflash assay in HCT116 cells in the presence of LGK974 (10nM). Treatment with LGK974 attenuated FOXQ1-dependent Wnt activation, especially in the presence of exogenous R-spondin 3. Data show one representative of n = 2 independent experiments with biological triplicates. **B.** qPCR of relevant Wnt ligands for comparison of FOXQ1 with FOXB2. Data are displayed as fold change compared to empty vector control and show biological triplicates. **C.** TOPflash assay in HCT116 cells to test FOXQ1-dependent Wnt activity upon treatment with CHIR99021 (5μM). FOXQ1 strongly potentiated Wnt signalling in synergy with CHIR99021. Data show one representative of n = 3 independent experiments with biological triplicates. **D.** TOPflash assay in HCT116 cells to test FOXQ1-dependent Wnt activity upon treatment with XAV939 (5μM). Wnt activity was abrogated upon β-catenin de-stabilization by XAV939. Data show one representative of n = 3 independent experiments with biological triplicates. **E.** TOPflash assay in 293T cells lacking β-catenin. FOXQ1 did not activate the β-catenin/TCF reporter in the absence of β-catenin. However, FOXQ1 synergized with exogenous wild-type and constitutively active β-catenin S33Y to activate Wnt signalling. Data show one experiment with biological triplicates. Data information: Data are displayed as mean ± SD. Statistical significance was calculated by ANOVA with Tukey’s post-hoc test (A, D, E) or Welch’s t-test (B, C), and defined as * *P* < 0.05, \*\**P* < 0.01, \*\*\**P* < 0.001.

**Figure EV3.**
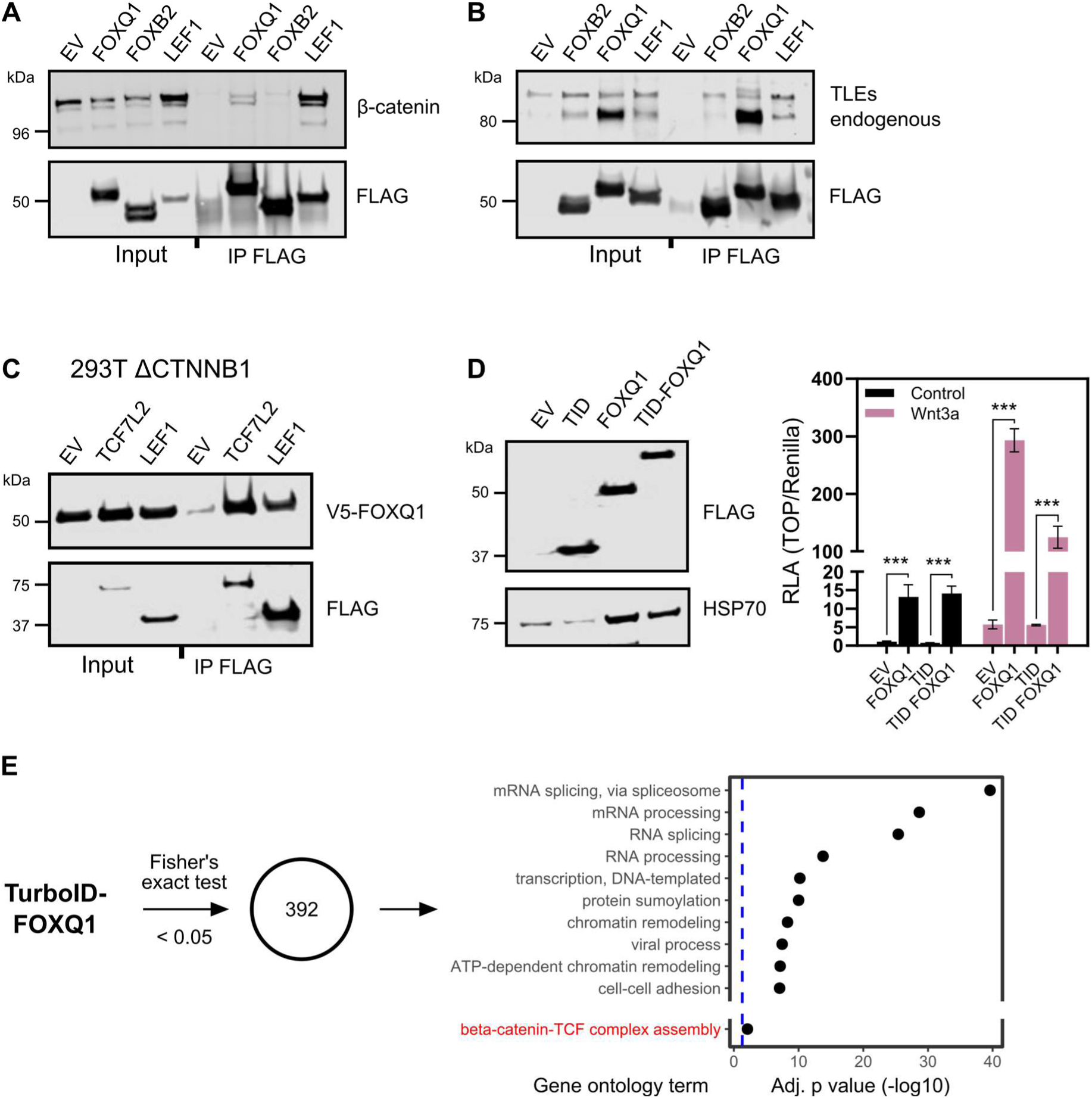
FOXQ1 interacts with β-catenin, TLEs, and TCF/LEF. **A.** Co-immunoprecipitation from nuclear lysates of 293T cells. Flag-tagged proteins were pulled down using Flag antibodies, and exogenous β-catenin mCherry was detected by immunoblot. FOXB2 and LEF1 were used as negative and positive controls, respectively. **B.** Co-immunoprecipitation from nuclear lysates of 293T cells. Flagged-tagged proteins were pulled down, and detection of endogenous TLEs was performed by immunoblot. **C.** Co-immunoprecipitation from nuclear lysates of 293T ΔCTNNB1 cells. Flag-tagged TCF7L2 and LEF1 proteins were pulled down in the presence of V5-FOXQ1. Immunoblot detection revealed TCF7L2/LEF1 interaction with FOXQ1 in the absence of β-catenin. A representative blot from n = 2 independent experiments is shown. **D.** Left: Immunoblot for protein expression of Flag-tagged TurboID-FOXQ1 fusion construct in 293T cells. TID, TurboID. Right: TOPflash assay for functional validation of the TID-FOXQ1 construct in 293T cells. The TID-FOXQ1 construct activated Wnt signalling similarly to wild-type FOXQ1. **E.** Gene ontology (GO) analysis of the statistically significant FOXQ1 interactors identified using TurboID. Only the top 10 most significant GO terms are shown in addition to *beta-catenin-TCF complex assembly*. Full results can be found in Table 1. The dashed blue line indicates an adjusted p-value of 0.05. Data information: Data are displayed as mean ± SD. Statistical significance was calculated by ANOVA with Tukey post-hoc test (D) and defined as * *P* < 0.05, \*\**P* < 0.01, \*\*\**P* < 0.001.

**Figure EV4.**
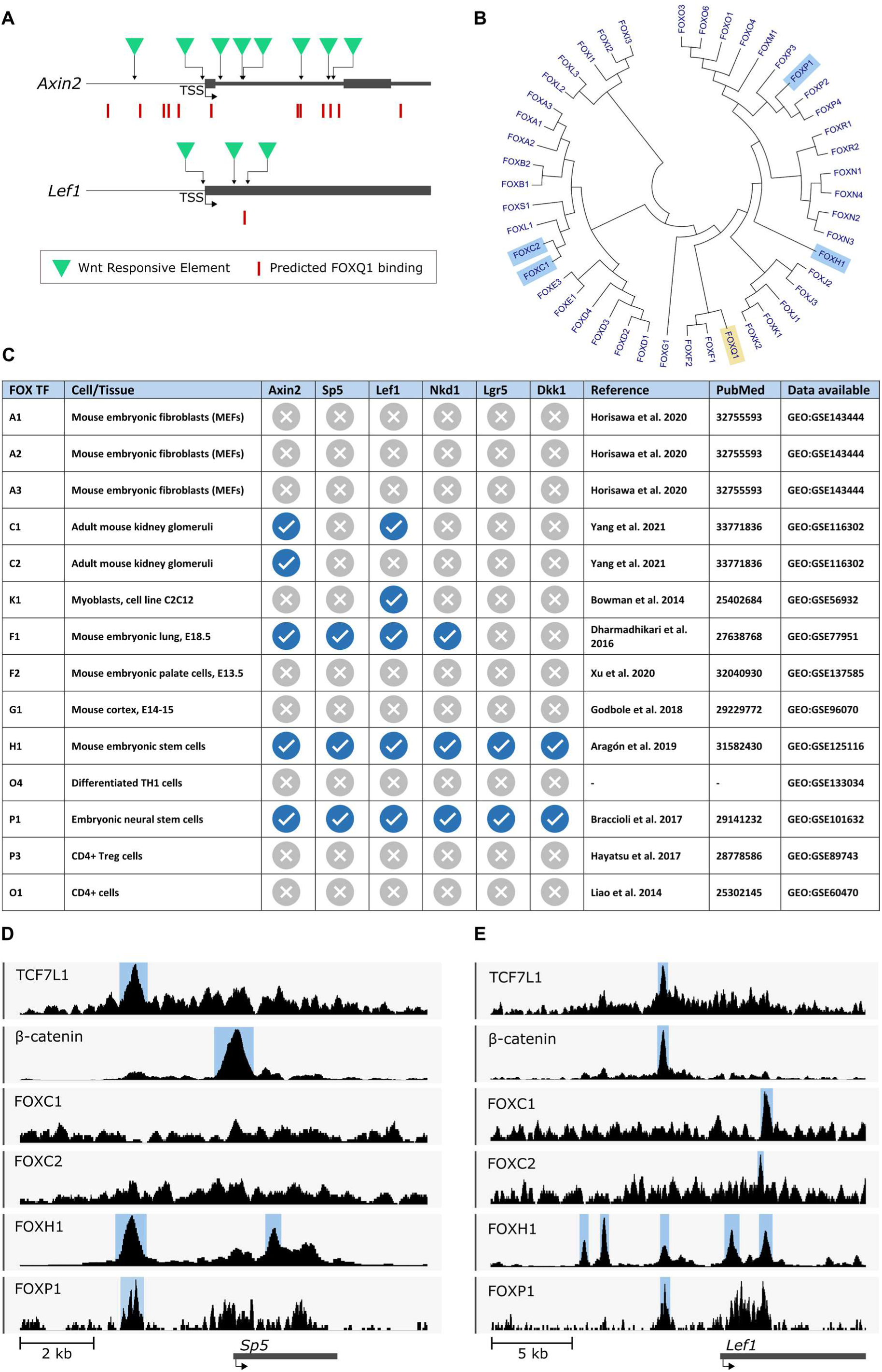
FOX transcription factors bind at known Wnt target genes. **A.** Prediction of FOXQ1 binding sites at the mouse *Axin2* and *Lef1* loci, based on JASPAR 2022 binding profile data. Green triangles denote Wnt-responsive elements (WREs) required for TCF/LEF binding, as previously identified (Jho *et al*., 2002; Li *et al*., 2006). Red rectangles denote predicted FOXQ1 binding sites. **B.** Phylogenetic tree of FOX transcription factors. Phylogenetic relationship between factors were determined based on their Forkhead box sequences. Highlighted in blue are FOX factors for which ChIP-seq genomic tracks are displayed. **C.** Table of FOX transcription factors for which ChIP-seq data has been obtained, including references to dataset depositories and associated publications. Blue check symbol denotes the presence of a called ChIP-seq peak at the promoter region of the corresponding gene. Gray cross symbol denotes the absence of a binding event. Note: for FOXH1 and FOXP1, only sequence coverage data (i.e., no peak calling data) were found, and the presence of binding events at gene promoters was assessed by visual inspection of these signalling tracks. **D.** Genomic tracks showing protein-DNA binding enrichment of TCF7L1, β-catenin, FOXC1, FOXC2, FOXH1 and FOXP1 at the *Sp5* locus, obtained from publicly available ChIP-seq datasets. **E.** Genomic tracks showing protein-DNA binding enrichment of TCF7L1, β-catenin, FOXC1, FOXC2, FOXH1 and FOXP1 at the *Lef1* locus, obtained from publicly available ChIP-seq datasets.

**Figure EV5.**
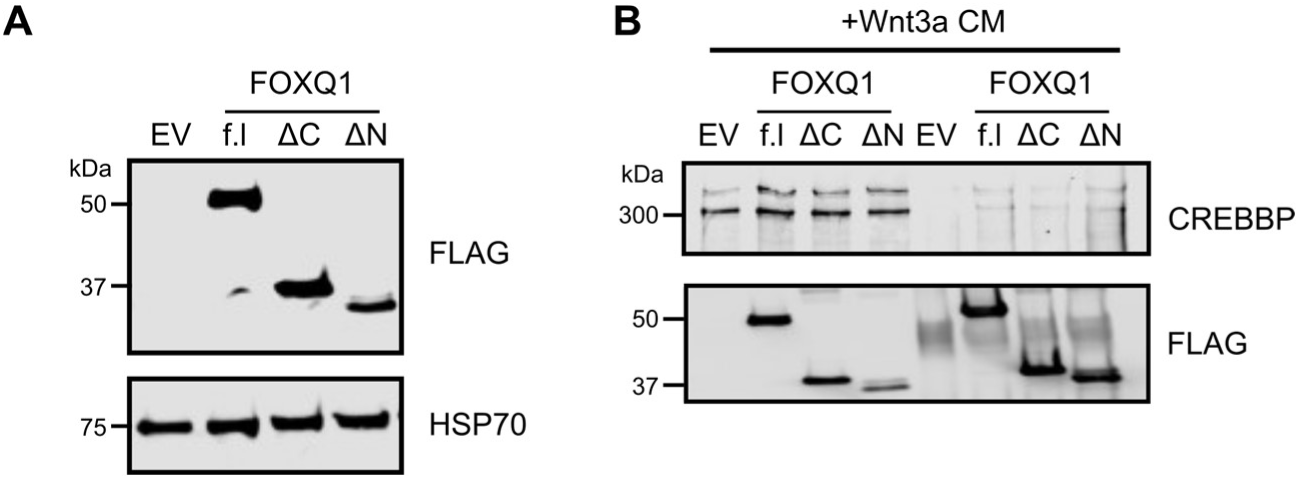
Distinct FOXQ1 protein domains differentially regulate Wnt signalling. **A.** Immunoblot of Flag-tagged FOXQ1 constructs in 293T cells. f.l: full-length. **B.** Co-immunoprecipitation from nuclear lysates of 293T cells. Flag-tagged FOXQ1 constructs were pulled down and endogenous CREBBP protein was detected by immunoblot. Cells were treated with Wnt3a conditioned media. All FOXQ1 constructs interacted with CREBBP.

## Tables Legends

**Table 1.** Gene Ontology (GO) analysis of FOXQ1 TurboID experiments

**Table 2.** SAINTexpress analysis of FOXQ1 TurboID experiments

## References

Bagati A, Bianchi-Smiraglia A, Moparthy S, Kolesnikova K, Fink EE, Lipchick BC, Kolesnikova M, Jowdy P, Polechetti A, Mahpour A et al. (2017) Melanoma Suppressor Functions of the Carcinoma Oncogene FOXQ1. Cell Rep 20: 2820–2832

Barker N, Hurlstone A, Musisi H, Miles A, Bienz M, Clevers H (2001) The chromatin remodelling factor Brg-1 interacts with beta-catenin to promote target gene activation. The EMBO journal 20: 4935–4943

Bourgeois B, Gui T, Hoogeboom D, Hocking HG, Richter G, Spreitzer E, Viertler M, Richter K, Madl T, Burgering BMT (2021) Multiple regulatory intrinsically disordered motifs control FOXO4 transcription factor binding and function. Cell Reports 36: 109446

Branon TC, Bosch JA, Sanchez AD, Udeshi ND, Svinkina T, Carr SA, Feldman JL, Perrimon N, Ting AY (2018) Efficient proximity labeling in living cells and organisms with TurboID. Nat Biotechnol 36: 880–887

Castro-Mondragon JA, Riudavets-Puig R, Rauluseviciute I, Berhanu Lemma R, Turchi L, Blanc-Mathieu R, Lucas J, Boddie P, Khan A, Manosalva Perez N et al. (2021) JASPAR 2022: the 9th release of the open-access database of transcription factor binding profiles. Nucleic Acids Res

Chavez A, Scheiman J, Vora S, Pruitt BW, Tuttle M, E PRI, Lin S, Kiani S, Guzman CD, Wiegand DJ et al. (2015) Highly efficient Cas9-mediated transcriptional programming. Nat Methods 12: 326–328

Chen H-C, Zhu Y-T, Chen S-Y, Tseng SCG (2012) Wnt signaling induces epithelial–mesenchymal transition with proliferation in ARPE-19 cells upon loss of contact inhibition. Laboratory Investigation 92: 676–687

Cho KF, Branon TC, Udeshi ND, Myers SA, Carr SA, Ting AY (2020) Proximity labeling in mammalian cells with TurboID and split-TurboID. Nature Protocols 15: 3971–3999

Christensen J, Bentz S, Sengstag T, Shastri VP, Anderle P (2013) FOXQ1, a novel target of the Wnt pathway and a new marker for activation of Wnt signaling in solid tumors. PLoS One 8: e60051

Clevers H (2006) Wnt/beta-catenin signaling in development and disease. Cell 127: 469–480

Dai S, Qu L, Li J, Chen Y (2021) Toward a mechanistic understanding of DNA binding by forkhead transcription factors and its perturbation by pathogenic mutations. Nucleic Acids Research 49: 10235–10249

Davidson G, Wu W, Shen J, Bilic J, Fenger U, Stannek P, Glinka A, Niehrs C (2005) Casein kinase 1 gamma couples Wnt receptor activation to cytoplasmic signal transduction. Nature 438: 867–872

Doumpas N, Lampart F, Robinson MD, Lentini A, Nestor CE, Cantu C, Basler K (2019) TCF/LEF dependent and independent transcriptional regulation of Wnt/beta-catenin target genes. EMBO J 38

Durinck S, Spellman PT, Birney E, Huber W (2009) Mapping identifiers for the integration of genomic datasets with the R/Bioconductor package biomaRt. Nature protocols 4: 1184–1191

Essers MAG, Vries-Smits LMMd, Barker N, Polderman PE, Burgering BMT, Korswagen HC (2005) Functional Interaction Between β-Catenin and FOXO in Oxidative Stress Signaling. Science 308: 1181–1184

Hoggatt AM, Kriegel AM, Smith AF, Herring BP (2000) Hepatocyte Nuclear Factor-3 Homologue 1 (HFH-1) Represses Transcription of Smooth Muscle-specific Genes *. Journal of Biological Chemistry 275: 31162–31170

Jho EH, Zhang T, Domon C, Joo CK, Freund JN, Costantini F (2002) Wnt/beta-catenin/Tcf signaling induces the transcription of Axin2, a negative regulator of the signaling pathway. Mol Cell Biol 22: 1172–1183

Jiang Y-G, Luo Y, He D-l, Li X, Zhang L-l, Peng T, Li M-C, Lin Y-H (2007) Role of Wnt/β-catenin signaling pathway in epithelial-mesenchymal transition of human prostate cancer induced by hypoxia-inducible factor-1α. International Journal of Urology 14: 1034–1039

Jiao X, Sherman BT, Huang DW, Stephens R, Baseler MW, Lane HC, Lempicki RA (2012) DAVID-WS: a stateful web service to facilitate gene/protein list analysis. Bioinformatics 28: 1805–1806

Kaneda H, Arao T, Tanaka K, Tamura D, Aomatsu K, Kudo K, Sakai K, De Velasco MA, Matsumoto K, Fujita Y et al. (2010) FOXQ1 is overexpressed in colorectal cancer and enhances tumorigenicity and tumor growth. Cancer Res 70: 2053–2063

Kirsch N, Chang L-S, Koch S, Glinka A, Dolde C, Colozza G, Benitez MD, De Robertis EM, Niehrs C (2017) Angiopoietin-like 4 is a Wnt signaling antagonist that promotes LRP6 turnover. Developmental cell 43: 71–82. e76

Kleinstiver BP, Prew MS, Tsai SQ, Topkar VV, Nguyen NT, Zheng Z, Gonzales APW, Li Z, Peterson RT, Yeh J-RJ et al. (2015) Engineered CRISPR-Cas9 nucleases with altered PAM specificities. Nature 523: 481–485

Knight JDR, Choi H, Gupta GD, Pelletier L, Raught B, Nesvizhskii AI, Gingras A-C (2017) ProHits-viz: a suite of web tools for visualizing interaction proteomics data. Nature methods 14: 645–646

Koch S (2021) Regulation of Wnt Signaling by FOX Transcription Factors in Cancer. Cancers (Basel) 13

Kolligs FT, Hu G, Dang CV, Fearon ER (1999) Neoplastic transformation of RK3E by mutant beta-catenin requires deregulation of Tcf/Lef transcription but not activation of c-myc expression. Molecular and cellular biology 19: 5696–5706

Kulak O, Chen H, Holohan B, Wu X, He H, Borek D, Otwinowski Z, Yamaguchi K, Garofalo LA, Ma Z et al. (2015) Disruption of Wnt/beta-Catenin Signaling and Telomeric Shortening Are Inextricable Consequences of Tankyrase Inhibition in Human Cells. Mol Cell Biol 35: 2425–2435

Li J, Sutter C, Parker DS, Blauwkamp T, Fang M, Cadigan KM (2007) CBP/p300 are bimodal regulators of Wnt signaling. The EMBO journal 26: 2284–2294

Li TW, Ting JH, Yokoyama NN, Bernstein A, van de Wetering M, Waterman ML (2006) Wnt activation and alternative promoter repression of LEF1 in colon cancer. Mol Cell Biol 26: 5284–5299

Liu J, Pan S, Hsieh MH, Ng N, Sun F, Wang T, Kasibhatla S, Schuller AG, Li AG, Cheng D et al. (2013) Targeting Wnt-driven cancer through the inhibition of Porcupine by LGK974. Proc Natl Acad Sci U S A 110: 20224–20229

MacDonald BT, Tamai K, He X (2009) Wnt/beta-catenin signaling: components, mechanisms, and diseases. Dev Cell 17: 9–26

Mellacheruvu D, Wright Z, Couzens AL, Lambert J-P, St-Denis NA, Li T, Miteva YV, Hauri S, Sardiu ME, Low TY et al. (2013) The CRAPome: a contaminant repository for affinity purification–mass spectrometry data. Nature Methods 10: 730–736

Moparthi L, Koch S (2020) A uniform expression library for the exploration of FOX transcription factor biology. Differentiation 115: 30–36

Moparthi L, Pizzolato G, Koch S (2019) Wnt activator FOXB2 drives the neuroendocrine differentiation of prostate cancer. Proc Natl Acad Sci U S A 116: 22189–22195

Moreira S, Seo C, Gordon V, Xing S, Wu R, Polena E, Fung V, Ng D, Wong CJ, Larsen B et al. (2018) Endogenous BioID elucidates TCF7L1 interactome modulation upon GSK-3 inhibition in mouse ESCs. bioRxiv

Nakayama J, Tan L, Li Y, Goh BC, Wang S, Makinoshima H, Gong Z (2021) A chemical screen based on an interruption of zebrafish gastrulation identifies the HTR2C inhibitor Pizotifen as a suppressor of EMT-mediated metastasis. eLife 10: e70151

Naujok O, Lentes J, Diekmann U, Davenport C, Lenzen S (2014) Cytotoxicity and activation of the Wnt/beta-catenin pathway in mouse embryonic stem cells treated with four GSK3 inhibitors. BMC Research Notes 7: 273

Nusse R, Clevers H (2017) Wnt/beta-Catenin Signaling, Disease, and Emerging Therapeutic Modalities. Cell 169: 985–999

Ohkawara B, Glinka A, Niehrs C (2011) Rspo3 binds syndecan 4 and induces Wnt/PCP signaling via clathrin-mediated endocytosis to promote morphogenesis. Dev Cell 20: 303–314

Ou C-Y, Kim JH, Yang CK, Stallcup MR (2009) Requirement of cell cycle and apoptosis regulator 1 for target gene activation by Wnt and beta-catenin and for anchorage-independent growth of human colon carcinoma cells. The Journal of biological chemistry 284: 20629–20637

Peng X, Luo Z, Kang Q, Deng D, Wang Q, Peng H, Wang S, Wei Z (2015) FOXQ1 mediates the crosstalk between TGF-beta and Wnt signaling pathways in the progression of colorectal cancer. Cancer Biol Ther 16: 1099–1109

Qiao Y, Jiang X, Lee ST, Karuturi RK, Hooi SC, Yu Q (2011) FOXQ1 regulates epithelial-mesenchymal transition in human cancers. Cancer Res 71: 3076–3086

Ramakrishnan AB, Chen L, Burby PE, Cadigan KM (2021) Wnt target enhancer regulation by a CDX/TCF transcription factor collective and a novel DNA motif. Nucleic Acids Res 49: 8625–8641

Robinson JT, Thorvaldsdóttir H, Winckler W, Guttman M, Lander ES, Getz G, Mesirov JP (2011) Integrative genomics viewer. Nature Biotechnology 29: 24–26

Slaymaker IM, Gao L, Zetsche B, Scott DA, Yan WX, Zhang F (2016) Rationally engineered Cas9 nucleases with improved specificity. Science 351: 84–88

Söderholm S, Cantù C (2020) The WNT/beta-catenin dependent transcription: A tissue-specific business. Wiley Interdiscip Rev Syst Biol Med: e1511

Tamura K, Stecher G, Kumar S (2021) MEGA11: Molecular Evolutionary Genetics Analysis Version 11. Mol Biol Evol 38: 3022–3027

Tan G, Lenhard B (2016) TFBSTools: an R/bioconductor package for transcription factor binding site analysis. Bioinformatics 32: 1555–1556

Tenbaum SP, Ordóñez-Morán P, Puig I, Chicote I, Arqués O, Landolfi S, Fernández Y, Herance JR, Gispert JD, Mendizabal L et al. (2012) β-catenin confers resistance to PI3K and AKT inhibitors and subverts FOXO3a to promote metastasis in colon cancer. Nature Medicine 18: 892–901

Teo G, Liu G, Zhang J, Nesvizhskii AI, Gingras A-C, Choi H (2014) SAINTexpress: improvements and additional features in Significance Analysis of INTeractome software. J Proteomics 100: 37–43

UniProt Consortium (2021) UniProt: the universal protein knowledgebase in 2021. Nucleic acids research 49: D480–D489

Veeman MT, Slusarski DC, Kaykas A, Louie SH, Moon RT (2003) Zebrafish Prickle, a Modulator of Noncanonical Wnt/Fz Signaling, Regulates Gastrulation Movements. Current Biology 13: 680–685

Walker MP, Stopford CM, Cederlund M, Fang F, jahn C, Rabinowitz AD, Graham DM, Yan F, Deal AM, Fedoriw Y, et al. (2015) FOXP1 potentiates Wnt/b-catenin signaling in diffuse large B cell lymphoma. Science Signaling 8: ra12

Xiang L, Zheng J, Zhang M, Ai T, Cai B (2020) FOXQ1 promotes the osteogenic differentiation of bone mesenchymal stem cells via Wnt/beta-catenin signaling by binding with ANXA2. Stem Cell Res Ther 11: 403

Yaklichkin S, Vekker A, Stayrook S, Lewis M, Kessler DS (2007) Prevalence of the EH1 Groucho interaction motif in the metazoan Fox family of transcriptional regulators. BMC Genomics 8

Yeo NC, Chavez A, Lance-Byrne A, Chan Y, Menn D, Milanova D, Kuo CC, Guo X, Sharma S, Tung A et al. (2018) An enhanced CRISPR repressor for targeted mammalian gene regulation. Nat Methods 15: 611–616

Zhang N, Wei P, Gong A, Chiu WT, Lee HT, Colman H, Huang H, Xue J, Liu M, Wang Y et al. (2011) FoxM1 promotes beta-catenin nuclear localization and controls Wnt target-gene expression and glioma tumorigenesis. Cancer Cell 20: 427–442

Zheng X, Lin J, Wu H, Mo Z, Lian Y, Wang P, Hu Z, Gao Z, Peng L, Xie C (2019) Forkhead box (FOX) G1 promotes hepatocellular carcinoma epithelial-Mesenchymal transition by activating Wnt signal through forming T-cell factor-4/Beta-catenin/FOXG1 complex. Journal of experimental & clinical cancer research 38: 1–15

